# Predicting the time needed to conduct an environmental systematic review or systematic map: analysis and decision support tool

**DOI:** 10.1101/303073

**Authors:** Neal R Haddaway, Martin J Westgate

## Abstract

Systematic reviews aim to maximise transparency and comprehensiveness, whilst also minimising subjectivity and sources of bias. Because of these time-consuming and complex tasks, systematic reviews are perceived as being resource-intensive. To date, published estimates of systematic review resource requirements have been largely anecdotal, being imprecise and not based on evidence. However, it is valuable to provide reliable means of estimating the resource and time requirements of systematic reviews and maps. We analysed all CEE systematic reviews (n=66) and maps (n=20) published or registered between 2012 and 2017 to estimate the average time needed to complete a systematic review and map. We then surveyed 33 experienced systematic reviewers to collate information on time needed for each stage of the review process. Our results show that the average CEE systematic review takes 157 days (SD; ±22), whilst the average CEE systematic map takes 209 days (SD; ±53). While screening of titles and abstracts is widely accepted to be time-consuming, in practice meta-data extraction and critical appraisal can take as long (or even longer) to complete, especially when producing systematic maps. Finally, we present a tool that allows the user to predict the time requirements of a review or map given information known about the planned methods and evidence base likely to be identified. Our tool uses evidence-based defaults as a useful starting point for those wishing to predict the time requirements for a particular review. Our analyses shed light on the most time-consuming stages of the systematic review and map process, and highlight key bottlenecks from the perspective of time requirements, helping future reviewers to plan their time accordingly. Future predictions of effort required to complete systematic reviews and maps could be improved if CEE and CEE review authors provided more detailed reporting of the methods and results of their reviewing processes.

## Introduction

Systematic review methods were developed in the field of healthcare in the 1990s as a means of collating, appraising and synthesising broad (and sometimes contradictory) bodies of primary research studies [1]. The methods revolve around a suite of practices during the conduct of a literature review that aim to maximise transparency and comprehensiveness, whilst also minimising subjectivity and sources of bias [2, 3]. Systematic reviews are now viewed as a ‘gold standard’ in evidence synthesis across not only healthcare [1], but also social welfare, education, international development, crime and justice [4], and conservation and environmental management [2]. Within these fields, not-for-profit organisations have been established to govern standards in systematic review and publish and endorse reviews that meet specific minimum standards (e.g. the Collaboration for Environmental Evidence, the Campbell Collaboration and Cochrane). Since their inception and development, the number of systematic reviews published by these coordinating bodies and more broadly across the research literature has increased considerably [3].

Systematic reviews should involve a number of important methodological steps to ensure the syntheses are reliable [5]. These include: 1) the publication of a peer-reviewed *a priori* protocol that sets out the planned methodology for the review, including detailed information regarding the search, screening, critical appraisal and data synthesis strategies; 2) comprehensive, tried-and-tested searches across a suite of resources for both traditional academic research studies and grey literature [6]; 3) screening of identified studies at title, abstract and full text levels using inclusion criteria that have been trialled and tested for consistency amongst reviewers; 4) considered critical appraisal of all sources of uncertainty and bias (validity) in each study, along with an assessment of the validity of all evidence collectively; 5) consistent extraction of data (both descriptive information, or meta-data, and quantitative or qualitative study findings); 6) accurate and reliable synthesis of study findings through appropriate quantitative (e.g. meta-analysis) or qualitative (e.g. meta-ethnography) methods; 7) throughout the process, full transparent documentation of all activities to allow verification and repeatability. Because of these time-consuming and complex tasks, systematic reviews are widely perceived as being particularly resource-intensive [7].

Although it is accepted that systematic reviews are challenging, the published estimates of the resource requirements of systematic reviews have been largely anecdotal, and as such are both imprecise and highly variable [e.g. 8]. One exception was a recent study by Borah et al. [9] who found that the average time from the date of registry to date of submission of final reports in the PROSPERO database was 67 weeks. There is notable uncertainty in this estimate, however, since the dates held by the PROSPERO database do not necessarily closely relate to the dates work commenced. Nor is there a clear link between the total duration of a systematic review and the actual time requirements in person-days. No comparable analysis of systematic review effort has been completed in the field of conservation and environmental management; but any such estimate based on protocol and final review report submission dates for the CEE journal Environmental Evidence (i.e. duplicating the approach of Borah et al.) is unlikely to be reliable. Indeed, an assessment of this data for the 86 reviews published by CEE between May 2012 and March 2017 suggest a mean time from protocol to review submission of 737 days (SD=±364) with a range of 48 to 1,524 days. At the lower range, this represents an impossible speed for review conduct, and at the upper end we know these data represent projects that underwent numerous significant hiatuses.

Since systematic reviews are known to be resource-intensive, and since current estimates of their time requirements are largely based on anecdote or uncertain data linked to reviews in other fields, there is a clear need to provide evidence-based estimates of the time needed to conduct a systematic review. Here, we report the results of a project that aimed to collate data from a variety of sources and summarise the time requirements of CEE systematic reviews. We use a combination of data reported within published systematic reviews and protocols along with data from a survey of systematic review practitioners in the environmental field. We produce an estimate of the mean time required to conduct a CEE systematic review or systematic map split by the key steps of the review process. We also describe a tool based on this data that allows those planning a systematic review or map to predict the time needed for their review based on their own scoping activities that reveal the likely volume of relevant evidence and the working speeds of their team. To our knowledge this is the first evidence-based tool for predicting workloads in a systematic review and has a broad applicability across a range of disciplines.

## Methods

### Assessment of published CEE SRs/SMs

An assessment was conducted of all CEE systematic reviews published since May 2012 in both the journal *Environmental Evidence*

(https://environmentalevidencejournal.biomedcentral.com/) and the CEE Library (http://www.environmentalevidence.org/completed-reviews). Key meta-data were extracted from all completed systematic reviews and systematic maps, along with systematic review and map protocols where no final review report had yet been published at the time of analysis (March 2017). This meta-data included the following: protocol submission date and review submission date (for all completed reviews and maps); the number of databases searched; the number of grey literature resources searched; the number of search results identified from database searching; the number of duplicates removed; the number of titles included after screening; the number of abstracts included after screening; the number of titles and abstracts included (where screened together); the number of full texts retrieved; the number of full texts included after screening; the number of studies included following critical appraisal; and, the number of studies with meta-analysable data. Data were separated according to whether they came from systematic maps or systematic reviews and summary figures and calculations were undertaken independently for these two types of review.

### Survey of systematic review practitioners

A list of potential respondents (n=61) was assembled from authorship lists of CEE systematic reviews, maps and protocols published between May 2012 and March 2017. The list was supplemented with personal contacts from the systematic review community (n=34). A total of 12 email addresses were no longer functioning and alternative authors from each EEJ publication were selected as target respondents. One further email address failed to work and so a tertiary alternative was selected and emailed. In total 95 respondents were targeted using functioning email addresses. An email invitation to an online survey was sent to each potential respondent (see Additional File 1 for survey questions). The key data collected are outlined in Table 1.

**Table 1.**
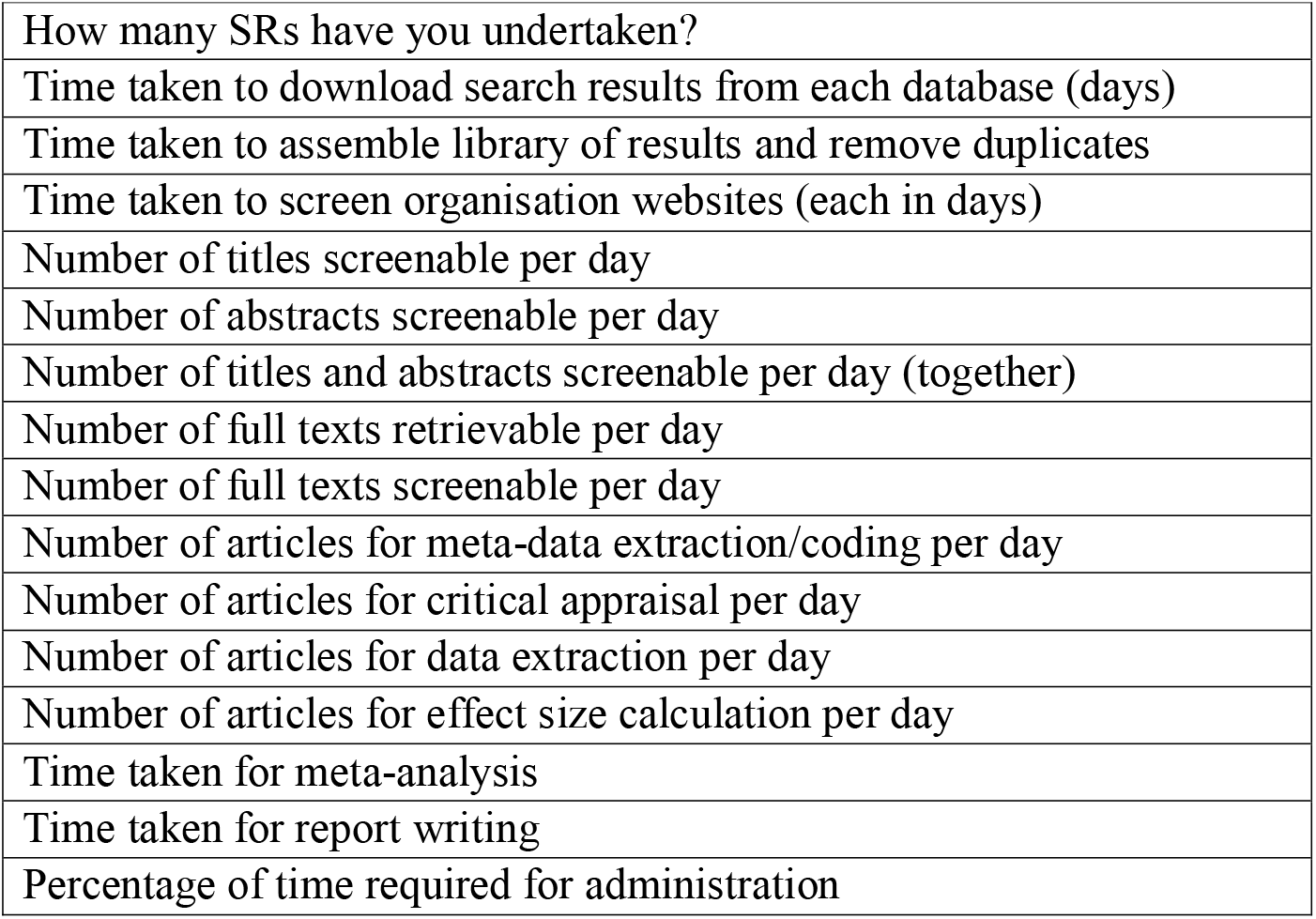
Key data collated from the survey of systematic review practitioners.

Some 30 responses were received through the online survey, yielding a response rate of 32%. Three responses were discarded because of incomplete information (only one page of responses received), resulting in a total of 27 valid responses. In addition, data from 6 systematic reviewers at one organisation in Canada were collated by their line manager and forwarded for use in the analysis. This resulted in a maximum of 33 data points or each question.

### Compilation of data and calculation of metrics

Following collation of the data from published articles and survey respondents, data were summarised using means and standard errors. Data were then transformed into the same units and information regarding the volume of evidence at each stage of the review process were combined with data on processing speeds to yield a set of summary data on the mean time taken for each main stage of the review process, along with a standard error. Standard errors were propagated for each individual calculation using an online error propagation tool (https://www.eoas.ubc.ca/courses/eosc252/error-propagation-calculator-fj.htm). The main stages of the review process were identified as outlined in Table 2. These stages are based on the CEE guidelines in systematic review [5]. Some data were arbitrarily set where CEE guidance exists (e.g. the percentage of titles used as a subset for testing consistency before commencing screening) or where data depend heavily on the experience level and efficiency of the reviewer (e.g. time taken for meta-analysis). Details of the sources of each line of data used in the calculation of times is provided in Table 3. Default values, the summary data and the calculations used to arrive at the metrics in Table 2 are provided in Additional File 2.

**Table 2.**
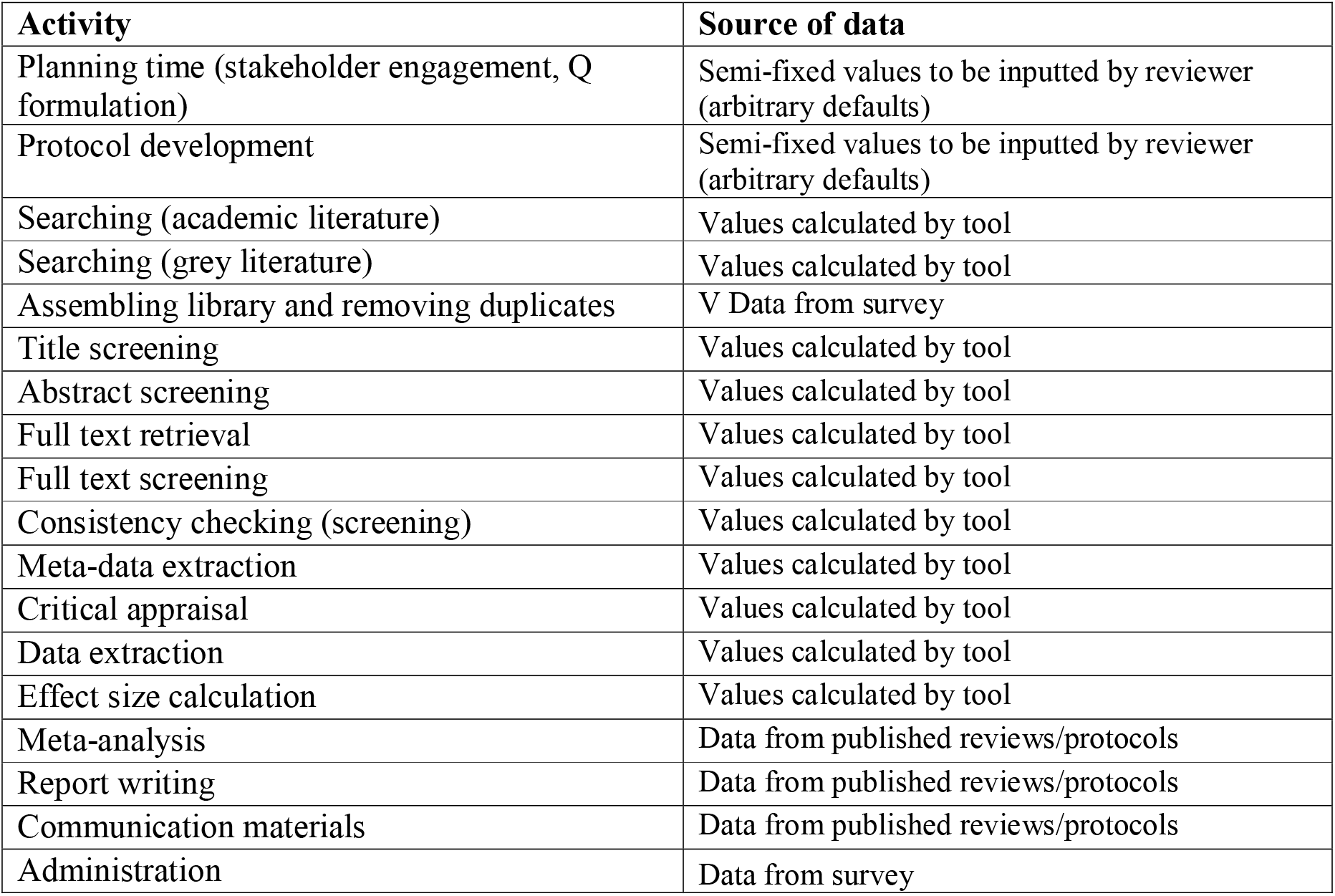
The main stages of a systematic review used to predict time requirements.

**Table 3.** Summary metrics used in intermediate steps of time calculation and the sources of data used to calculate metrics in Table 2.

| Activity | Source of data |
| --- | --- |
| Number of searching databases | Data from published reviews/protocols |
| Time taken to download one database's search results (days) | Data from survey |
| Number of grey literature website sources | Data from published reviews/protocols |
| Time required for additional grey literature sources (other languages, etc.) (days) | Semi-fixed values to be inputted by reviewer (arbitrary defaults) |
| Number of grey literature sources (website) screenable per day | Data from survey |
| Number of search results (total) | Data from published reviews/protocols |
| Estimated duplicate rate (percent) | Data from published reviews/protocols |
| Number of search results after duplicate removal | Data from published reviews/protocols |
| Estimated inclusion rate (title) (percent) | Data from published reviews/protocols |
| Screening speed (titles per day) | Data from survey |
| Estimated inclusion rate (abstract) (percent) | Data from published reviews/protocols |
| Screening speed (abstracts per day) | Data from survey |
| Retrieval rate (percent retrievable) | Data from published reviews/protocols |
| Retrieval speed (articles per day) | Data from survey |
| Estimated inclusion rate (full text) (percent) | Data from published reviews/protocols |
| Screening speed (full texts per day) | Data from survey |
| Percentage of results screened for consistency (title) | Semi-fixed values to be inputted by reviewer (arbitrary defaults) |
| Percentage of results screened for consistency (abstract) | Semi-fixed values to be inputted by reviewer (arbitrary defaults) |
| Percentage of results screened for consistency (full text) | Semi-fixed values to be inputted by reviewer (arbitrary defaults) |
| Number of consistency checkers at each stage | Semi-fixed values to be inputted by reviewer (arbitrary defaults) |
| Meta-data extraction rate (articles per day) | Data from survey |
| Critical appraisal rate (articles per day) | Data from survey |
| Percentage of studies with meta-analysable data | Data from published reviews/protocols |
| Data extraction rate (articles per day) | Data from survey |
| Effect size calculation rate (articles per day) | Data from survey |
| Estimated inclusion rate following critical appraisal | Data from published reviews/protocols |
| Percentage of time needed for administration | Data from survey |

### A software tool for estimating effort in future reviews

Following calculation of summary time metrics for each stage of the review process (and propagated standard errors), an interactive research effort estimation tool was produced that builds on our framework by allowing end users to replace the default or mean data with specific values based on their own experiences or knowledge. Key requirements for the tool were: transparency in indicating the sources and evidence behind default values through methodological documentation provided herein; helping the user understand the nature of each step in the review process; building in details and instructions from published guidance on systematic reviews; and, ease of use. The aim of this tool is to provide an indication of the minimum time requirements for a systematic review. It is hoped that this tool will continue to develop as the dataset upon which it is based expands and the models are refined. The structure that provides caveats for missed steps is therefore intended as a conservative warning where it is not based on evidence, and is informed by existing published systematic review methodology and guidance.

The tool is provided in two formats. One format is based in an Excel spreadsheet, since this format is downloadable and readily usable. The second format is a web-based app, which is more easily updated and refined. The app was built in the R statistical environment [10] using the R packages Shiny [11] and shinydashboard [12] to construct the interactive framework, and plotly [13] to draw the diagrams. Both software tools make identical calculations, and return identical results.

The software tool utilises several different types of user input. First, it requires an initial number of articles that are returned by the ‘search’ stage of the systematic review or systematic map. This is typically easy to estimate, as it is simply the sum of the number of hits from all databases searched during the review. The software tool then combines this total number of articles with estimates of the proportion of articles retained at each stage (i.e., title screening, abstract screening, etc.), and the rate at which articles can be processed during those stages. Typically, the proportion of articles retained increase as the review progresses, while the number processed per day decreases. Finally, the user can add estimates of the time taken to undertake specific tasks within the review process, such as conducting a metaanalysis or writing a report. These data are then combined into plots of the number of articles expected, and the total time spent, on each review stage.

The tool underwent substantial revisions and alterations during our analyses. Over time, we developed an increasing level of detail to reflect the variability and nuance across the suite of activities that make up a systematic review or map. Our final tool is published here along with detailed explanatory notes to guide the user through its use and to ensure that reliable, contextualised data (i.e. through scoping) is provided where possible to increase the accuracy of predictions. The excel version of the app is available in the supplementary information (Additional File 3), while the web app can be used online at https://mjwestgate.shinyapps.io/revtime/ or downloaded for use in R using the source code on github (https://github.com/mjwestgate/revtime/).

## Results

### Assessment of published CEE SRs

A total of 108 systematic review publications were produced by CEE between May 2012 and March 2017, of which 66 represented systematic reviews and 20 were systematic maps (86 in total), though 35 of these documents (41%) were protocols for as-yet incomplete projects. The majority of the data comes from systematic reviews, and of these data the majority relate to as yet unfinished systematic reviews (Figure 1).

**Figure 1.**
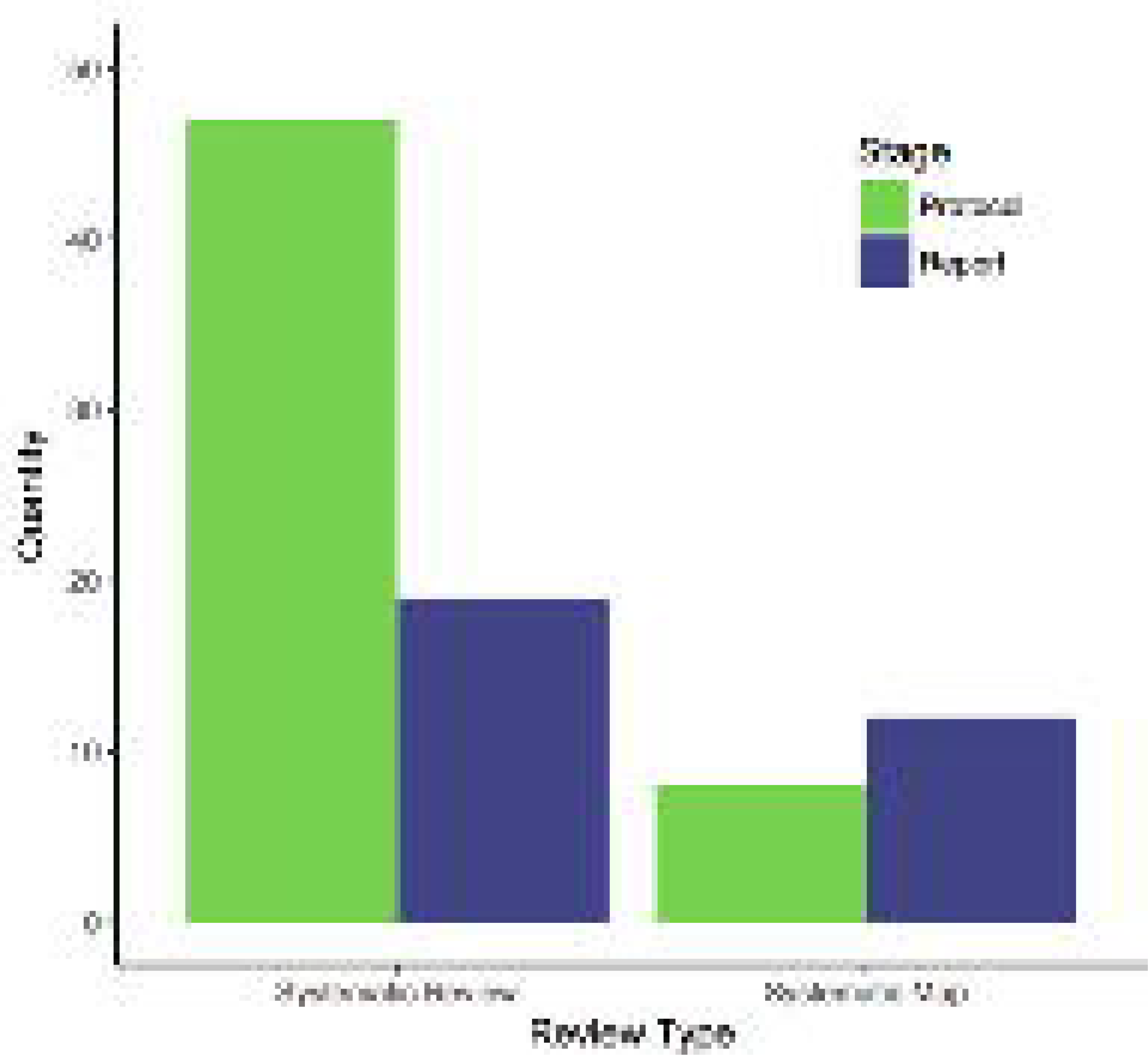
The number of publications from CEE between May 2012 and March 2017.

The mean number of records remaining after each key stage in the conduct of a systematic review is outlined in Figure 2 and Table 4. The variability around the data is clearly large, particularly for some points in the review process where data were lacking (e.g. meta analysis, n=3). Some notable reviews could be perceived as outliers: for example, the systematic review on the timing of mowing impacts on biodiversity in meadowland that resulted in a particularly small set of search results (n=367) and a relatively high inclusion rate at title screening stage (74.0%) [14], and the systematic map of on-farm water quality mitigation measures that resulted in a very large set of search results (n> 145,000) and a relatively high percentage of duplicates (49.5%) [15]. However, given the low sample size these cases have been left in to reflect the real variability present.

It is also worth noting that there is a lack of consistent reporting in published systematic reviews and maps. Despite the existence of published standards for the reporting of activities in systematic reviews (e.g. PRISMA; [16]) and requirements for a high level of detail in reporting within the journal *Environmental Evidence*, only 8 or the 32 completed systematic reviews and maps reported data for all stages of the review process (i.e. searching, duplicate removal, title, abstract and full text screening and full text retrieval).

**Figure 2.**
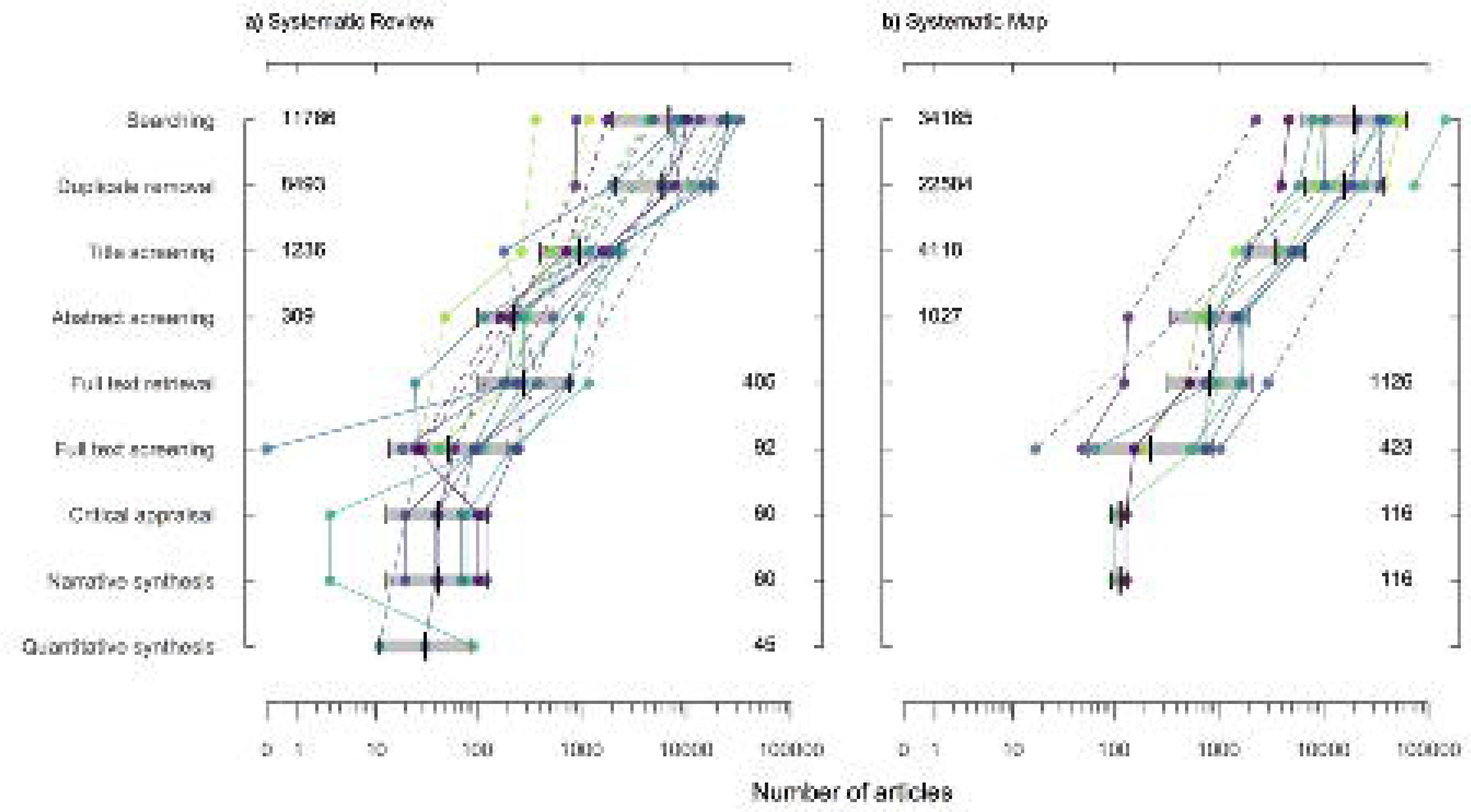
The mean number of records remaining after each key stage of the a) systematic review and b) systematic map processes. Error bars are ±1 standard deviation. Labels are data values.

**Table 4.** Mean number of records remaining in each stage of the systematic review and map process, along with minima, maxima, standard errors (SE) and sample sizes (n=number of reviews/protocols providing data).

| Number of items remaining after: | Mean | SD | SE | n<sup>543</sup> |
| --- | --- | --- | --- | --- |
| Searching | 11,786 | 10,230 | 2347 | 19 |
| Duplicate removal | 8,493 | 6,040 | 2135 | 8 |
| Title screening | 1,236 | 766 | 231 | 11 |
| Abstract screening | 309 | 269 | 85 | 10 |
| Full text retrieval | 405* | 347 | 105 | 11 |
| Full text screening | 93 | 85 | 20 | 18 |
| Critical appraisal | 60 | 41 | 15 | 8 |
| Narrative synthesis | 60 | 41 | 15 | 8 |
| Quantitative synthesis | 45 | 43 | 25 | 3 |

*Systematic maps*
| Number of items remaining after: | Mean | SD | SE | n<sup>546</sup> |
| --- | --- | --- | --- | --- |
| Searching | 34,165 | 39,434 | 11,384 | 12 |
| Duplicate removal | 22,584 | 20,578 | 6,204 | 11 |
| Title screening | 4,118 | 2,002 | 817 | 6 |
| Abstract screening | 1,027 | 575 | 217 | 7 |
| Full text retrieval | 1,126* | 859 | 286 | 9 |
| Full text screening | 423 | 371 | 117 | 10 |
| Critical appraisal | 116 | 22 | 16 | 2 |
| Narrative synthesis | 116 | 22 | 16 | 2 |

\* The number of full texts is greater than the number of screened abstracts due to the inclusion of grey literature at the full text stage that have been externally screened for relevance from the searching of websites, web-based search engines, etc.

### Survey of systematic review practitioners

Of the 33 included responses, only 7 provided data for all 15 questions asked about their experience with reviews, while a further 12 provided data for the stages up to data/meta-data extraction and beyond. On average, respondents had conducted a median of 2 systematic reviews (minimum=0, maximum=18). Only one respondent had not previously conducted a review before: data from this respondent were in relation to full text retrieval alone, since they had acted as an assistant for a larger group of reviewers. We received fewer responses about later stages of the review than early stages (Table 5), and particularly few responses about the time taken to complete quantitative synthesis (effect size calculation and meta analysis; n=7 and 8 respectively).

**Table 5.** Summary data for respondents to the survey: n, sample size; SD, standard deviation.

| Survey question | Mean response | n | SD |
| --- | --- | --- | --- |
| Time taken to download search results from each database (days) | 0.25 | 20 | 0.220 |
| Time taken to assemble library of results and remove duplicates | 1.37 | 18 | 1.403 |
| Time taken to screen organisation websites (each in days) | 0.15 | 21 | 0.122 |
| Number of titles screenable per day | 854.35 | 23 | 533.622 |
| Number of abstracts screenable per day | 192.29 | 24 | 111.900 |
| Number of titles and abstracts screenable per day (together) | 468.14 | 22 | 128.216 |
| Number of full texts retrievable per day | 170.94 | 24 | 137.369 |
| Number of full texts screenable per day | 43.99 | 30 | 31.014 |
| Number of articles for meta-data extraction/coding per day | 16.69 | 21 | 11.574 |
| Number of articles for critical appraisal per day | 11.68 | 19 | 8.145 |
| Number of articles for data extraction per day | 6.87 | 19 | 5.093 |
| Number of articles for effect size calculation per day | 24.00 | 7 | 34.083 |
| Time taken for meta-analysis | 6.75 | 8 | 5.092 |
| Time taken for report writing | 15.53 | 20 | 10.228 |
| Percentage of time required for administration | 19.00 | 22 | 12.276 |

A typical CEE systematic review results in a mean of just over 11,000 search results, which falls to approximately 8,500 unique records following duplicate removal (Table 4). Just over 1,200 records remain following title screening and around 300 following abstract screening. With the addition of evidence from other sources, the total number of full texts obtained is on average c. 400. Screening of these full texts leaves just over 90 relevant articles/studies. Critical appraisal retains approximately 60 articles/studies, and suitable data are present in a little over 40 of them.

The sample size for systematic maps was much smaller than for systematic reviews (n=20 versus n=66), but the volume of evidence was far greater for these maps: almost 35,000 search results were obtained on average, leaving over 20,000 unique records. Title screening left over 4,000 relevant records and abstract screening left over 1,000. Just over 1,100 full texts were retrieved, with over 400 being relevant at full text. Across the two cases where critical appraisal was performed within a systematic map, on average of over 100 studies were retained in the final map.

### Estimated effort

The time taken for each stage of a systematic review were lower, on average, for the corresponding stage of a systematic map (Figure 3). The total time estimated for an ‘average’ systematic review is 157 days (SD; ±22), whilst the total time for an ‘average’ systematic map is 252 days (SD; ±67) when including an optional critical appraisal step, or 209 days (SD; ±53) excluding critical appraisal. This estimate includes a large amount of time allotted to planning and administration, in an effort to be conservative (45 days for systematic reviews and 60 days for systematic maps [including critical appraisal]). Stages that are calculated by the model include those from searching to effect size calculation, whilst other stages are set as arbitrary defaults that must be changed by the user (see ‘The tool’, below). For these calculated stages, the most time consuming are title screening, full text screening and critical appraisal, with meta-data and data extraction also requiring considerable time. Searching (for traditional academic and grey literature), assembling a library of evidence, full text retrieval, and consistency checking required less time than most other stages. The uncertainty around this data is substantial, resulting from the propagation of errors across the models and the variability in the underlying source data.

**Figure 3.**
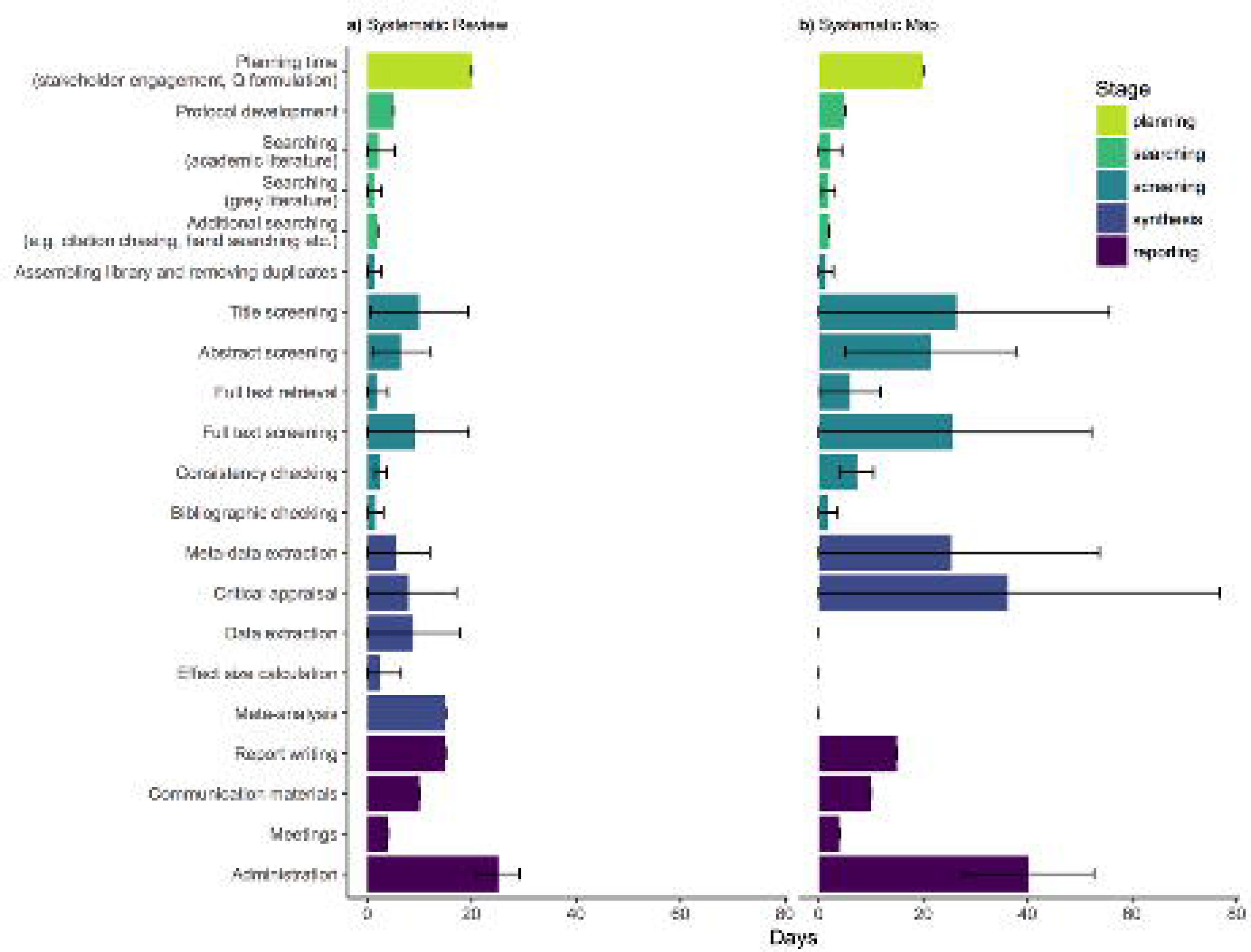
Time taken for each stage of the systematic map process in days. Error bars are ±1 standard deviation.

## Discussion

In this paper, we have presented the most comprehensive estimate to date of the effort needed to complete environmental systematic reviews and systematic maps. Our results revealed substantial variability in the number of articles included in synthesis projects, and the total time taken to complete them. However, we also found key bottlenecks during early screening and at critical appraisal and data extraction stages of each review. Below we expand on these findings and their implications for future research practice.

### Emergent patterns

We found several trends during our analysis that did not match our expectations about which stages of the review would take the most time. Particularly surprising was the observation that a relatively small proportion of time is spent on performing searching activities: between 7 and 7.5 days for reviews and maps, respectively. This result may reflect detailed preparation, given that searching should be preceded by in depth building and testing of search strategies, that will be outlined in an *a priori* protocol. Whilst we did not explicitly ask expert reviewers how long they spent designing and testing a search strategy, this part of the review process requires careful planning to ensure the review results are comprehensive and representative of the true evidence base for a particular topic [17, 18].

Equally unexpected was our finding that respondents’ reported time spent on administration was particularly large: on average 19% of their total time. For reviews this corresponded to 24 days, whilst for maps it was almost 40 for those including critical appraisal (35.5 for those without critical appraisal). Reported administration time varied substantially (SD=12.3), perhaps indicating discrepancies in respondents’ definitions of what should be included. However, this likely reflects the fact that systematic reviewing often requires time spent coordinating a large, possibly international team, and may also require substantial learning or ‘relearning’ of particular skills, such as experimental design or statistics. We have not factored in training time in our analysis, but this is worth considering for novel teams or those that will rely heavily on group tasks using subject but not methodology expertise.

More expected was the large amount of time spent on screening (including retrieval); an average of 80 and 27 days for systematic reviews and maps respectively. This is a large proportion of the time budget (17% and 32% for reviews and maps, respectively [39% for maps excluding critical appraisal]). These differences highlight the fact that resources are predominantly shifted towards identifying evidence in maps, whereas far more time is devoted to synthesis in reviews. In reviews a similar time is spent on extracting and analysing the data as screening (25 days). In maps, however, the proportion of total time on extracting meta-data and coding is relatively lower (also 25 days).

### The implications for ‘optional’ activities

Our calculations allow us to estimate the impacts of various optional activities on the total time requirements of a systematic review or map. Current CEE guidance suggests that a subset of articles is checked for consistency in the application of inclusion criteria between two reviewers prior to commencing screening in earnest [5], and it suggests that 10% of records should be checked as a minimum. However, in the field of healthcare systematic reviews, dual coding is common [e.g. 19]. By altering the level of consistency checking from the recommended minimum of 10% at each stage to 100% (i.e. complete dual screening), the total time required changes from 155 days to 183 days, an increase of 18%. Whilst regarded by some as a gold standard for systematic review methodology [1], this increase in time requirements is substantial and may prove too costly for some. However, it may be an important concession to maximise reliability and minimise human error in some cases.

Similarly, the CEE guidance suggests a selection of review bibliographies is screened to help to maximise comprehensiveness of the search [5]. Increasing this bibliographic checking or ‘citation chasing’ can require considerable time if, for example, all identified reviews are screened in this way, or even if all articles’ bibliographies are screened. Assuming that the inclusion rate at title, abstract and full text (and retrieval rate) remain the same in bibliographic checking as for the core of the review, one can readily predict the additional time needed to screen a certain number of reviews or articles in this way. Within a systematic map, a larger volume of reviews is likely to be found, and the user can specify this number.

For example, in a systematic map of the impacts of vegetated strips within and around fields [20], around 100 review bibliographies were checked for additional potentially relevant articles. Altering the number of bibliographies checked in our tool to 100 increases the time requirement from 255 to 271 days (6%).

### Comparison with existing estimates

Previous estimates of the resource requirements of systematic reviews have been imprecise. and vary substantially, from between 6 months and 24 months or several years (Table 6). Anecdotally, we have heard estimates that are as long as 5 years by a leading institute that produces systematic reviews in healthcare in Sweden (SBU,http://www.sbu.se/en/). Our analyses demonstrate that the time requirements for an ‘average’ CEE-style systematic review need only take 157 days (FTE). This estimate represents just under 1 year FTE, taking vacation, public holidays, and other regular disruptions to full time work into account. Therefore, our analysis reveals a resource requirement in the lower end of the rough estimates provided in the literature. Interestingly, our estimate is under half that of the only other evidence-based assessment of which we are aware, which corresponds to approximately 337 days [9]. It is vital to remember that the time estimate by Borah et al. (2017) and the other rough time estimates in the literature are typically meant to reflect the time required to conduct a systematic review, rather than the resource requirements. However, the average total salary costs for a postdoctoral research at Bangor University (chosen arbitrarily due to our knowledge of the university, including National Insurance and USS pension contributions) for 12 months is 48,593 GBP at the time of writing (https://www.bangor.ac.uk/finance/py/documents/pay-scales-en.pdf). Including other costs, such as support staff time and travel and subsistence for meetings, this sum is unlikely to rise above 100,000 GBP. This value, again, sits below the mid points for the roughly estimated cost ranges provided in the literature.

**Table 6.** Estimates of the resource requirements for systematic reviews from a non-systematic search of the literature.

| Financial cost | Time requirement | Review type | Citation |
| --- | --- | --- | --- |
|  | 0.5-3 years | Systematic review | Dicks, L.V., Walsh, J.C. and Sutherland, W.J., 2014. Organising evidence for environmental management decisions: a '4S' hierarchy. *Trends in ecology & evolution*, 29(11), pp.607-613. |
| 30,000-300,000 USD | Several years | Systematic review | Collaboration for Environmental Evidence. 2013. Guidelines for Systematic Review and Evidence Synthesis in Environmental Management. Version 4.2. Environmental Evidence: www.environmentalevidence.org/Documents/Guidelines/Guidelines4.2.pdf |
| 80,000-120,000 GBP | 10-18 months | Systematic review | Collins, A., Coughlin, D., Miller, J. and Kirk, S., 2015. The production of quick scoping reviews and rapid evidence assessments: a how to guide. |
| <=250,000 USD |  | Systematic review | McGowan, J. and Sampson, M., 2005. Systematic reviews need systematic searchers. *Journal of the Medical Library Association*, 93(1), p.74. |
|  | 9-24 months | Systematic review | Centre for Cognitive Ageing and Cognitive Epidemiology (http://www.ccace.ed.ac.uk/research/software-resources/systematic-reviews-and-meta-analyses) |
|  | 67.3 weeks | Systematic review | Borah, R., Brown, A.W., Capers, P.L. and Kaiser, K.A., 2017. Analysis of the time and workers needed to conduct systematic reviews of medical interventions using data from the PROSPERO registry. *BMJ open*, 7(2), p.e012545. |

Our estimates do not attempt to predict a full costing of a systematic review. Furthermore, our analyses are based on reported volumes of work and efficiency rates from the literature and expert systematic reviewers. As such, the numbers are in need of validation using detailed, accurate records of recently completed reviews. However, our estimates and tool are a useful starting point for those wishing to better understand the demands and likely time requirements of a systematic review or map.

The times estimated by our tool for the time required to undertake a systematic review are realistic relative to the reviews conducted by Mistra EviEM. This 6 year project funded by Mistra to undertake systematic reviews and maps in the field of environment relevant to the Swedish environmental goals (www.eviem.se/en). The project will have completed 17 systematic reviews and maps over a 6.5 year period, with c. 20 years of full time equivalent staff resources (review project managers); approximately 2.2 years of time per review. However, our estimates for systematic maps are somewhat higher than those indicated for EviEM maps in our experience. This is almost certainly the result of a small and heterogeneous evidence base for completed systematic maps: fewer systematic maps have been completed to date and the variability around the volume of evidence is substantial (SD for systematic review total search results is 11,786 records, whist it is 39,434 for systematic maps). Systematic maps are more adaptable by their very nature [21, 22], but a larger evidence base would be useful in increasing the precision of the data in our tool.

### Limitations of our analysis and the evidence base

Our analysis presents the best available information on the number of articles, and the amount of time, included in a typical environmental systematic review or map. However, it is possible that a number of factors may adversely affect the reliability of our analyses of the ‘average’ time needed to complete a review or map. Below we outline some of these points so as to avoid the risk of faulty interpretation of our results.

First, our systematic review calculations assume a quantitative synthesis will be performed, which may be true for the majority of current systematic reviews in the field of environment. However, qualitative synthesis is a valuable evidence synthesis method [23], and its use will likely increase in CEE reviews in the future. However, qualitative systematic reviews often do not hold the same regard for issues such as comprehensiveness that quantitative systematic reviews do. For example, qualitative syntheses may stop screening after a certain point because of information saturation. Accordingly, these reviews should be dealt with different when performing an analysis relating to time requirements, and tools for predicting times should be built specifically for these kinds of analysis. It may be the case, however, that specific qualitative reviews could adapt our tools to fit the desired methods. Our intention, however, is not to make a universal analysis or tool for all types of reviews. Instead, we focus on traditional quantitative systematic reviews.

Second, all of the data in our analyses have a high level of variability, poor levels of reporting, or both. This results both from a heterogeneous evidence base and a relatively small sample size. For example, of the 19 completed systematic reviews, only 8 reported the number of duplicates removed from total search results. Similarly, whilst 18 reviews reported the total number of included articles, only 10-11 articles reported the number of articles following title screening, abstract screening and full text retrieval. Future CEE reviews should strive to report such methodological information consistently, and CEE should create or adhere to accepted reporting standards for all published reviews (e.g. PRISMA [16] or ROSES [24]). Future analyses should increase sample size, making the most of the rapidly expanding body of reliable systematic reviews. Indeed, efforts to record descriptive summary information regarding systematic review methods are underway (e.g. ROSES [24]).

Third, we were not able to provide evidence-based data for all parts of our analysis. A number of key variables have been estimated from our personal experience, including the time required for additional searching for literature, and the number of bibliographies screened. Where possible, future analyses should attempt to examine the evidence base for these data.

Fourth, there are clear exceptional circumstances that would affect the reliability of predictions made from our tool. For example, a change in core staff midway through a project would likely require a significant proportion of time to acquaint new staff with what has been done to date. However, careful file management and clear record keeping could reduce this to a minimum. Large review teams may require more resources to train and manage, particularly if meeting remotely. Novice teams may require substantial training time and may suffer from low efficiency in the earlier stages of a review. Finally, undertaking reviews over an extended time period can result in particularly low efficiency if core staff must reacquaint themselves with their own work after significant gaps.

Fifth, our tool allows the end user to estimate the time required to complete a systematic review or systematic map, and our analyses of the evidence base provide useful default values should any information be unknown to the user. These default values, however are based on an ‘average’ systematic review or map. It is important to note that the heterogeneity across CEE reviews means that this ‘average’ review, whilst helpful as a starting point, is perhaps not a meaningful entity. Context is highly important for each review, and knowing something about the volume or the nature of the evidence (e.g. proportional relevance of a subset) will allow end users to estimate time requirements much more accurately. We should not assume that all reviews are alike and that the times calculated in our analysis are a reliable estimate alone when planning a review. We encourage users to undertake good quality scoping, as suggested in the CEE Guidelines [5] so as to provide reliable predictions of the volume of evidence, the proportional relevance of articles and studies, and the time required by the user’s team to undertake specific tasks.

Finally, we have calculated mean volumes of evidence at each stage of the review process and have used inclusion rates and working speeds to calculate an independent mean time requirement for each stage based on available evidence. However, many reviews do not report all data for each stage of the review, and the results of one stage are dependent upon the nature of the stages preceding it. In ideal circumstances, we would have full data from all reviews that would allow us to model the time requirement based on various contextual variables, for example the inclusion rate of the preceding stage. This is not possible with our limited dataset, however, and our methods represent a necessary compromise.

### Future work needed

As described above, there is a need for a greater number of data points in future analyses, both for published systematic reviews and maps and for survey data relating to processing speeds. This itself would be aided by better reporting of methods used and records found at all stages of the review process in CEE reviews. Some efforts are underway to record this data more consistently (e.g. ROSES [24]).

We also highlight the need for evidence-based estimates of the financial costs associated with systematic reviews, taking into account the price of necessary software, consultancy support (e.g. from an informatician), registration and publication fees, communication materials, and physical meetings. Although there will be considerable local and regional variability in the real world prices of these services, an itemised list of recommended activities is a vital point of departure for those planning an efficient and successful review.

Finally, results from our analyses and predictions using our tool should be continually tested and the tool refined in order to match developments in systematic review methodology (e.g. machine learning and prioritised screening [25]).

## Conclusions

Our analyses shed light on the most time-consuming stages of the systematic review and map process. We have highlighted key bottlenecks from the perspective of time requirements, and our results allow future reviewers to plan their time accordingly. Our tool uses evidence-based defaults as a useful starting point for those wishing to predict the time requirements for a particular review. We also call on CEE and CEE review authors to improve the reporting of the methods and results of their reviewing processes.

## Acknowledgements

The authors thank Sif Johansson and Mistra EviEM for contributing time for the completion of this manuscript.

## Additional Files

### Additional File 1

Email survey sent to systematic review practitioners.

### Additional File 2

Data and calculations used to arrive at metrics and standard errors compiled to produce time requirements for the various steps of a systematic review.

### Additional File 3

Systematic review and map time planner tool (Excel version).

